# Intact protein barcoding enables one-shot identification of CRISPRi strains and their metabolic state

**DOI:** 10.1101/2024.06.05.597362

**Authors:** Vanessa Pahl, Paul Lubrano, Daniel Petras, Hannes Link

**Author notes:** shared first authors (Vanessa Pahl, Paul Lubrano).

## Abstract

Simultaneous measurements of library barcodes and metabolites can facilitate the screening of metabolically engineered bacterial libraries. Here, we introduce a scalable method for the parallel measurement of protein-based library barcodes and metabolites using flow-injection mass spectrometry (FI-MS). Protein barcodes are based on ubiquitin, a small 76-amino-acid protein, which we labeled with N-terminal tags of six amino acids to identify bacterial strains. We demonstrate that FI-MS effectively detects intact ubiquitin proteins and identifies the N-terminal barcodes. In the same analysis, we semi-quantify relative concentrations of primary metabolites. As a proof of concept, we engineered six ubiquitin-barcoded CRISPRi strains targeting metabolic enzymes, and analyzed their metabolic profiles and ubiquitin barcodes using FI-MS. Our results show distinct metabolome changes within metabolic pathways targeted by CRISPRi. This method enables fast and simultaneously detection of library barcodes and intracellular metabolites, opening up new possibilities to measure metabolomes of engineered bacteria at scale.

## Main

Rapid identification of bacterial strains within genetic libraries is important for synthetic biology applications, especially for screening of synthetic metabolic pathways and engineered enzymes. Traditional methods involve DNA barcoding, where unique DNA sequences are integrated into each bacterial strain. These sequences are then amplified and matched against a barcode database using high-throughput sequencing technologies^1,2^. DNA barcoding offers high specificity but lacks the capability to capture molecular phenotypes of engineered bacteria in the same measurement, because DNA barcoding is limited to inference of fitness phenotypes via barcode frequency. Alternatively, RNA-barcodes can be visualized using fluorescence microscopy combined with fluorescent in situ hybridization (FISH) to detect more complex phenotypes and for in *situ* genotyping of CRISPR libraries^3^. Recently, CRISPR screens have been combined with time-of-flight mass cytometry (CyTOF) or immunohistochemistry for multiplexed cell barcoding at the protein level^4,5^. This method used protein barcodes (Pro-Codes) with triplet epitopes linked to specific guide RNAs for multidimensional protein profiling in single-cell pooled screens. Protein barcodes were also used for high throughput screens of protein binding libraries using so called flycodes and mass spectrometry^6^. Yet, there are currently no methods to simultaneously measure barcodes and metabolome. Such measurements could provide almost real-time strain identification, and can significantly reduce the time and cost required for arrayed CRISPR screens.

To address this, we introduce a novel method that uses high-throughput mass spectrometry to detect protein barcodes and metabolites in the same sample and in the same measurement. As a case study we used CRISPR interference (CRISPRi) in *E. coli* metabolism. For CRISPRi we used a catalytically dead Cas9 mutant (dCas9) to inhibit transcription of a target gene defined by the base pairing region of the single-guide RNA (sgRNA)^7^. To label CRISPRi strains with a protein barcode, we co-expressed the sgRNA and a small protein from the same plasmid (**Figure 1a**). As protein barcode we selected human ubiquitin, a small protein consisting of 76 amino acids, which can be detected by intact protein analysis via top-down mass spectrometry^8^. To create unique barcodes, we engineered ubiquitin by adding a six-amino acid tag to its N-terminal (**Figure 1a**). Initially, we added a random sequence - LVFYHA - to ubiquitin, which resulted in the LVFYHA-ubiquitin barcode. This barcode was expressed under a constitutive promoter from the same plasmid as the sgRNA. Ubiquitin and sgRNA were oriented in opposite directions, with a 38 bp spacer that separates their respective promoter sequences. This design enables the exchange of sgRNA protospacer sequences and N-terminal tags using a single 241 bp DNA oligonucleotide (**Figure 1a**).

**Figure 1.**
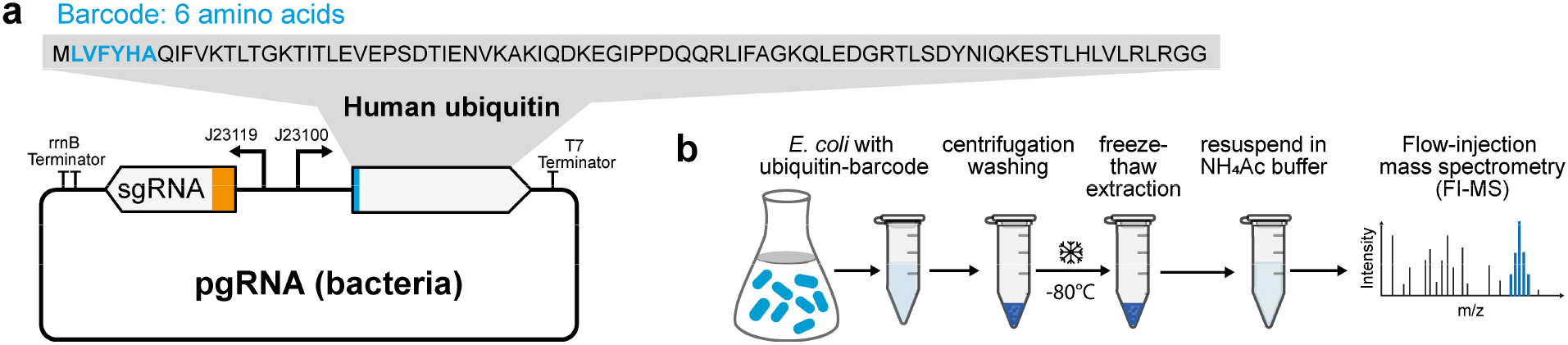
Protein barcoding with ubiquitin and detection by flow injection mass spectrometry. a) The gene encoding human ubiquitin was integrated next to the single guide RNA on the plasmid pgRNA bacteria^7^. A N-terminal sequence of 6 amino acids serves as a barcode. Shown is the LVFYHA-ubiquitin barcode. b) Sampling protocol for simultaneous detection of metabolites and intact ubiquitin by flow-injection mass spectrometry (FI-MS).

First, we confirmed expression of the LVFYHA-ubiquitin barcode in an *E. coli* CRISPRi strain, which carried a non-targeting sgRNA, and evaluated barcode detection by mass spectrometry. As a control strain, we used the same CRISPRi strain but without the ubiquitin barcode. Cell extracts were prepared from cultures in exponential growth phase using a single freeze-thaw cycle and resuspension in ammonium carbonate buffer to ensure intact protein recovery (**Figure 1b**). The cell extracts were then analyzed by flow-injection time-of-flight mass spectrometry (FI-MS)^9,10^. FI-MS detected 1044 m/z features that matched theoretical masses of the LVFYHA-ubiquitin barcode, considering different charge states, sodium adducts and 13-C isotopes (**Figure 2a**). Importantly, 73% of these features were significantly more abundant in the strain that expressed LVFYHA-ubiquitin compared to the control strain (p-value < 0.01 and fold-change > 4), thus demonstrating reproducible detection of the barcode across three replicates (**Figure 2b**). Most significant m/z features resulted from the LVFYHA-ubiquitin species that had 9 and 10 positive charges, which are within an m/z range of 930 to 1050 (**Figure 2c**). No significant features were detected in the lower mass range below 600, which is the mass range of most primary metabolites, and therefore we assumed that ubiquitin does not interfere with metabolite detection. Finally, we used the FLASHdeconv tool^11^ for spectral deconvolution of the FI-MS data. This analysis accurately identified the monoisotopic mass of LVFYHA-ubiquitin at 9290.00 Da (**Figure 2d**). In summary, these results demonstrate that ubiquitin with an N-terminal tag is expressed in *E. coli* and that this barcode can be effectively identified with FI-MS and a fast extraction protocol that does not require peptide digestion.

**Figure 2.**
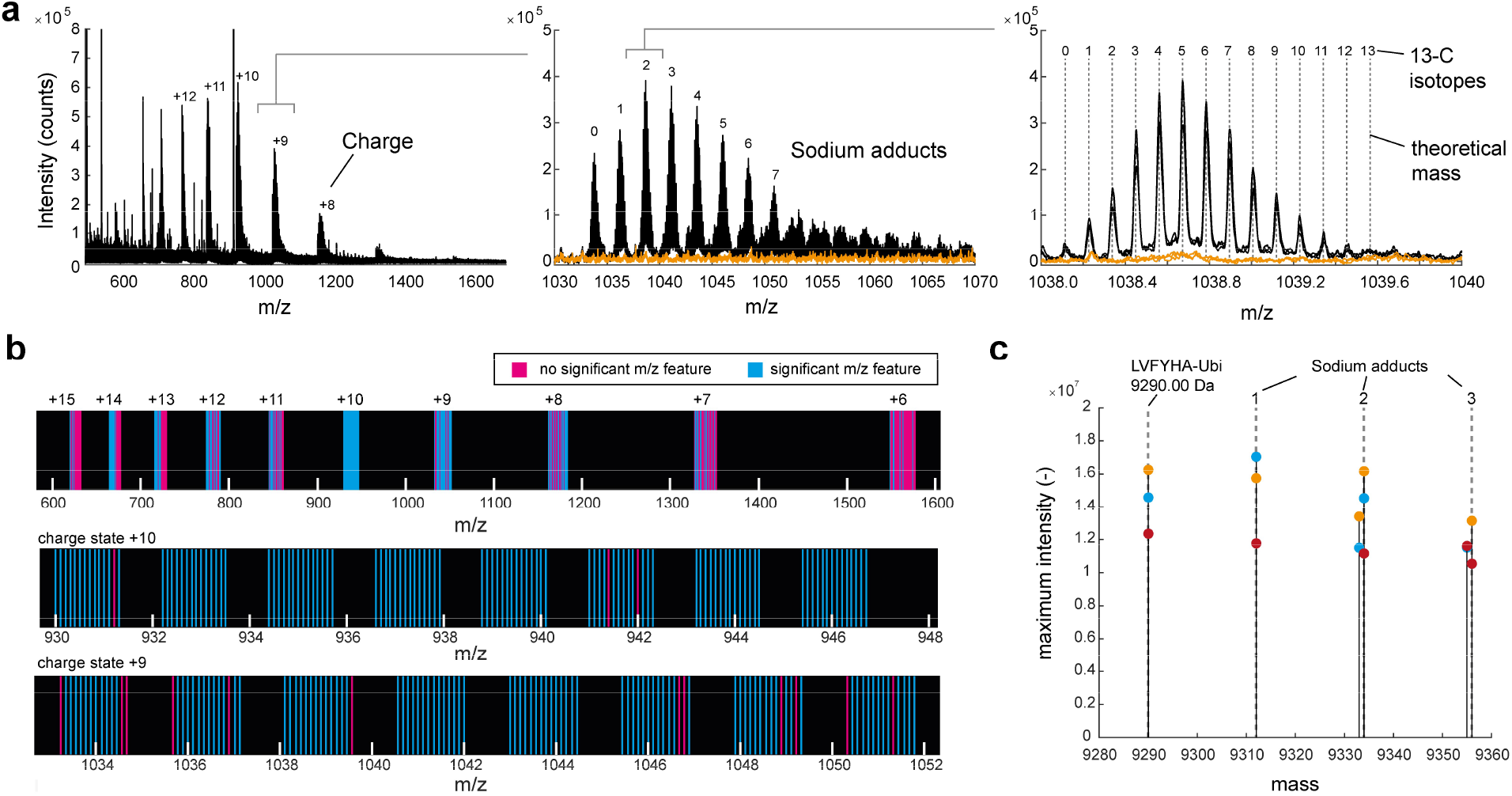
FI-MS detects the LVFYHA-Ubiquitin barcode. a) Mass spectra of cell extracts from *E. coli* expressing LVFYHA-ubiquitin (black). Extracts from a control strain without ubiquitin are shown in orange. Shown is the MS1 spectrum in positive mode from 500 to 1700 mass-to-charge ratio (m/z) measured with FI-MS. Different charge states are due to varying degrees of protonation, z = 8-12 are indicated (left). Sodium adducts of the +9 charge state, 0-7 sodium adducts are indicated (middle). 13-C isotope distribution of z =+9 charge state and 2 sodium adducts, exact monoisotopic masses are shown as dashed lines (right). b) m/z features were annotated to exact monoisotopic masses of LVFYHA-ubiquitin with different charges, sodium adducts and 13-C isotopes. Annotated m/z features that were significantly different in LVFYHA-ubiquitin samples relative to the control samples are shown in blue (n= 3 samples, p<0.01 and fold-change >4). c) Spectral deconvolution of MS1 spectra of LVFYHA-ubiquitin identified the exact monoisotopic mass of LVFYHA-ubiquitin (9290.00), and sodium adducts. Shown are 5 monoisotopic masses with highest summed intensity of n = 3 replicates. Dashed lines are exact masses and dots with different colour are deconvoluted masses of n = 3 replicates.

Next, we created six additional ubiquitin barcodes, each with a distinct mass, and tested if we can distinguish these barcodes based on their mass alone (**Figure 3a**). The six barcodes were combined with sgRNAs targeting different metabolic genes. Four of the target-genes encoded enzymes in biosynthesis pathways of the amino acids histidine (*hisD*), lysine (*dapE*), threonine (*thrC*) and leucine (*leuB*). Two other target-genes encoded isocitrate dehydrogenase (*icd*) in the tricarboxylic acid cycle, and 1-deoxy-D-xylulose 5-phosphate reductoisomerase (*dxr*) in the methylerythritol phosphate pathway. The control strain was again the LVFYHA-ubiquitin barcode in combination with a non-targeting sgRNA.

**Figure 3.**
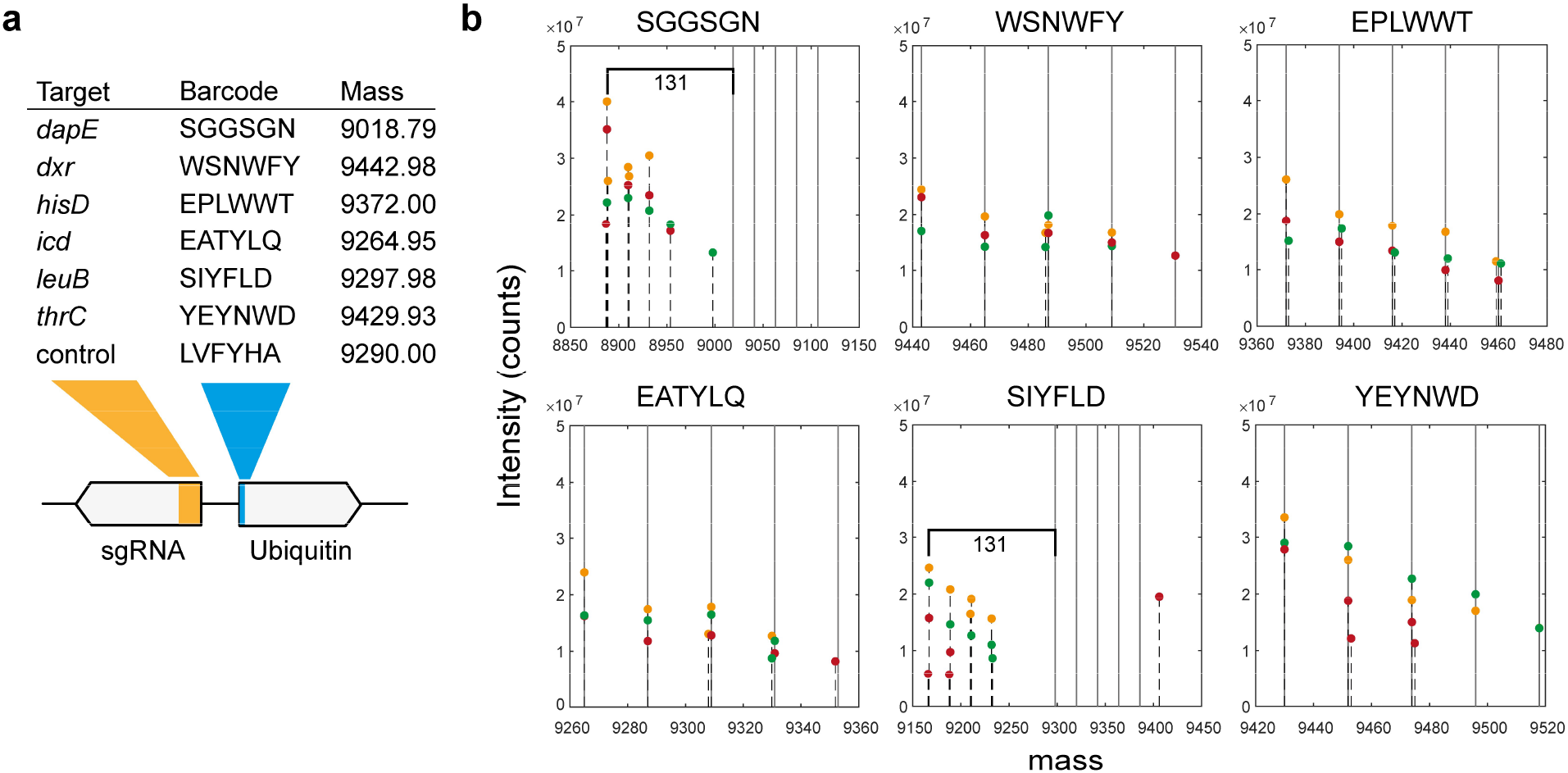
Ubiquitin barcodes of 6 CRISPRi strains and their spectral deconvolution. a) Table of CRISPRi strains. Shown is the gene targeted by the sgRNA and the 6 amino acids of their ubiquitin barcode. The monoisotopic mass of the ubiquitin barcode is given in Dalton. b) Monoisotopic masses determined by spectral deconvolution of FI-MS data. Shown are the 5 monoisotopic masses with highest summed intensity from n = 3 samples. Grey lines are the theoretical masses of the barcode with 0-4 sodium. Dots with different colour are deconvoluted masses of n = 3 replicates.

Metabolites and proteins of these strains were sampled after 6.5 hours of growth in minimal glucose medium and anydrotetracycline (aTc) to induce the CRISPRi system. FI-MS analysis of the extracts showed distinct monoisotopic masses for each barcode (**Figure 3b**). In four strains (*dxr, icd, hisD, thrC*) the 5 most abundant monoisotopic masses matched the theoretical monoisotopic mass of the respective barcode and up to four sodium adducts. Few signals determined by spectral deconvolution differed by 1 Da, presumably due to errors in assignment of either charge states or isotope distribution. In two strains (*dapE, leuB*) spectral deconvolution did not result in the correct monoisotopic mass of the respective barcodes (SIYFLD, SGGSGN). In both cases the mass difference between measured and theoretical mass was -131 Da, which corresponds to the mass of a peptide-bound methionine. This suggested hydrolytic cleavage of the N-terminal methionine from the SIYFLD-ubiquitin and SGGSGN-ubiquitin barcodes. In *E. coli*, the enzyme methionine aminopeptidase (MAP) facilitates the removal of the first methionine residue in proteins, and its activity is higher when the first amino acid has small side chains^12^, such as the serine (S) of the SIYFLD-ubiquitin and SGGSGN-ubiquitin barcodes. Consequently, the design of the ubiquitin barcodes could be optimized by selecting amino acids with larger side chains to avoid methionine cleavage.

Having established that FI-MS can detect the ubiquitin barcodes, we investigated if we can detect metabolites in the same FI-MS data. FI-MS detected 526 m/z features that we could annotate to 701 metabolites in the genome-scale metabolic model of *E. coli i*ML1515^13^. Principle component analysis separated the different CRISPRi strains (**Figure 4a**), thus indicating that our method is robust and detects distinct metabolic states induced by the CRISPRi perturbations. Next, we inspected metabolites in the metabolic pathways that are associated to the CRISPRi perturbation, which are biosynthesis pathways of histidine, leucine, threonine and lysine, the MEP pathway and the TCA cycle (**Figure 4b**). The *hisD* strain showed strong increases of 5 intermediates in the histidine pathways, including the *hisD* substrate metabolite histidinol (histd). This increase of histidine intermediates is likely caused by a bottleneck at the end of the pathway through the knockdown of *hisD*. Similarly, FI-MS detected an accumulation of substrate metabolites of the other three CRISPRi strains that target amino acids pathways: N-succinyl-L-2,6-diaminoheptanedioate (sl26da) in the *dapE* strain, O-phospho-L-homoserine (phom) in the *thrC* strain, and (2R,3S)-3-isopropylmalate (3c2hmp) in the *leuB* strain. Furthermore, the substrate metabolites of the *icd* and dxr strain, isocitrate (icit) and 1-deoxy-D-xylulose 5-phosphate (dxyl5p), increased respectively. These results show that our approach detects distinct metabolome changes in the CRISPRi strains, including stronger metabolome changes in the pathways targeted by CRISPRi (e.g. increase of substrate metabolites in all 6 CRISPRi strains).

**Figure 4.**
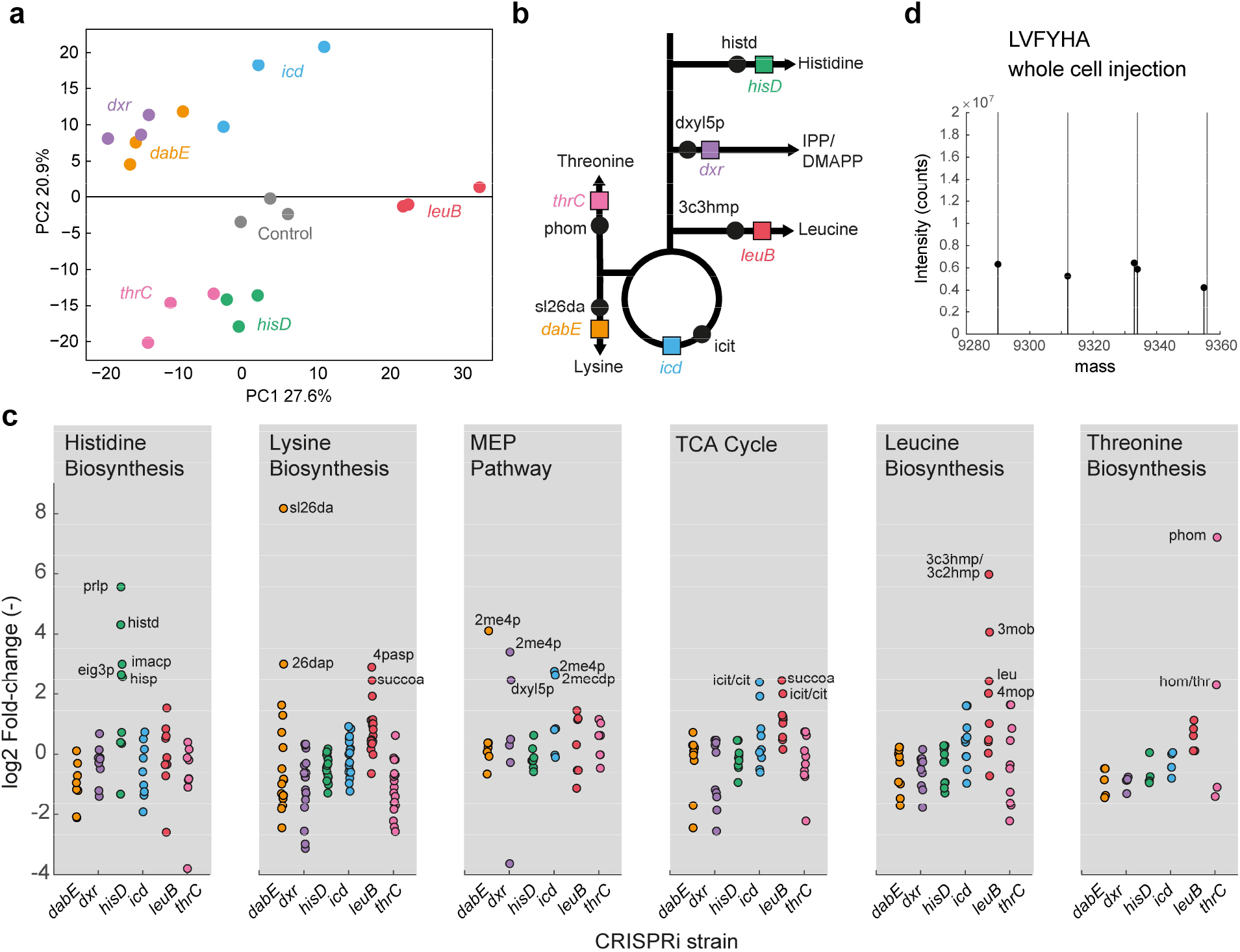
Ubiquitin-barcodes allows simultaneous analysis of metabolic signature of CRISPRi strains. a) Principle component analysis of m/z features that were annotated to metabolites. Shown are PC1 and PC2 of 701 metabolites measured in n= 3 samples of 7 CRISPRi strains. b) CRISPRi target pathways. Shown are the target enzymes and their substrate metabolites. c) Levels of metabolites in metabolic pathways that are targeted by the six CRISPRi strains. Metabolite levels are shown as log2 fold change relative to the control strain with a non-targeting sgRNA. Dots show the mean from n = 3 cultures. Metabolites are grouped by pathways, and metabolites with log2 FC > 3 are labelled. d) Analysis of the ubiquitin-LVFYHA barcode in whole living cells. The CRISPRi control strain expressing LVFHA-ubiquitin was diluted in 1:8 M9-medium: water and 3 μL were directly injected into the mass spectrometer. Cells were heated to 90°C in the heat exchanger of the column oven. Shown are 5 deconvoluted masses with highest summed intensities.

In conclusion, we introduced a novel approach for barcoding CRISPRi strains using ubiquitin, which is integrated with the sgRNA in a dual expression system. Our method combines aqueous extraction and MS1 spectral analysis, which efficiently captures both protein barcodes and primary metabolites. Apart from CRISPRi strains, the approach should enable rapid identification of other libraries of engineered bacteria and it has advantages over traditional methods that require more time-consuming and labor-intensive plasmid extraction and sequencing. The method could identify unique barcode signatures and concentration changes across hundreds of metabolites, within 1-minute analysis time per sample. It is also possible to skip the extraction step, as we identified the LVFYHA-ubiquitin barcode by injecting whole living cells directly into the mass spectrometer (**Figure 4d**). Thus, future studies could measure intact protein barcodes with real-time metabolomics^14^ or single-cell metabolomics^15^ to screen metabolic states of CRISPR libraries with ultra-high-throughput. Although we only generated a small set of protein tags for this feasibility study, it is important to point out that at MS/MS level it is possible to differentiate also isobaric tags, thus opening up the possibility to generate millions of potential sequence tags. Therefore, we anticipate that our method could be the starting point for further development of MS/MS level differentiation of sequence tags, and for ultra-high-throughput metabolome screening in libraries of engineered bacteria with broad applications in synthetic biology.

## Acknowledgement

This work was funded by the DFG Cluster of Excellence EXC2124 ‘Controlling Microbes to Fight Infection’ (CMFI).

## Authors contribution

Conceptualization: all authors; Methodology: all authors; Investigation: all authors; Visualization: VP, HL; Funding acquisition: DP, HL; Project administration: HL; Supervision: HL; Writing – original draft: VP, HL.

## Methods

### Media

Cultivations were performed with LB medium (Cat#L3522, Sigma-Aldrich) or M9 minimal medium with glucose (5 g/L) as sole carbon source. The M9 medium consists of (per liter): 7.52 g Na_2_HPO_4_·2 H_2_O, 5 g KH_2_PO_4_, 1.5 g (NH_4_)2SO_4_, 0.5 g NaCl. Additionally, the following components were sterile filtered and added separately (per liter M9 medium): 1 mL 0.1 M CaCl_2_, 1 mL 1 M MgSO_4_, 0.6 mL 0.1 M FeCl_3_, 2 mL 1.4 mM thiamine-HCl and 10 mL trace salts solution. The trace salts solution consists of (per liter): 180 mg ZnSO_4_·7 H2O, 120 mg CuCl_2_·2 H_2_O, 120 mg MnSO_4_·H_2_O, 180 mg CoCl_2_·6 H_2_O. For strains transformed with pgRNA-ubiquitin plasmid or variants, 100 μg/mL carbenicillin (Carb) was added to the media. To induce the expression the dCas9 protein, aTc was supplemented to a final concentration of 200 nM.

### Strains and culture

*E. coli* YYdCas9^3^ was the wild-type strain used in this study and all strains in this study derived from this strain. MegaX DH10B competent *E. coli* (Cat#C640003, Invitrogen) were used for cloning.

### Cloning of pgRNA-ubiquitin

The ubiquitin coding sequence, T7 terminator and RBS were first amplified from the plasmid Ubiquitin WT^16^ (Cat#12647, Addgene) and added to pgRNA-bacteria^7^ using Golden gate cloning^17^. The promotor J23100 was amplified from a gene fragment that was ordered from Twist Bioscience (San Francisco, USA) and was added to the pgRNA using IVA cloning^18^, forming pgRNA-ubiquitin. The plasmid was then transformed into *E. coli* YYdCas9 using TSS transformation^19^.

### Cloning of CRISPRi strains

Single stranded, 241 bp long DNA oligonucleotides were ordered from Twist Bioscience (San Francisco, USA). Oligos were amplified and added to pgRNA-ubiquitin to replace the mRFP protospacer and add a tag to the ubiquitin coding sequence. Gibson assembly^20^ was used and ligation products were transformed in *E. coli* MegaDH10B with electroporation. Colonies were picked and plasmids extracted using miniprep (Thermo Fisher #K0503). After sequencing of ubiquitin barcode and protospacer region, the plasmids were transformed in *E. coli* YYdCas9 using electroporation.

### Cultivation and sampling of ubiquitin-barcodes and metabolites

For metabolite and ubiquitin-barcode analysis, strains were inoculated from a glycerol stock in 3 mL rich medium (LB) with carbenicillin (100 μg/mL) and cultivated for 8 h at 37°C. Cells were then transferred in 10 mL M9 minimal medium with glucose and carbenicillin for overnight cultivation at 37°C in 100 mL shake flasks. M9-pre-cultures were adjusted to a starting OD_600_ of 0.05 into 10 mL M9 medium. The strains were cultivated for 6.5 h. Subsequently, OD_600_ was measured and an equivalent of 1 OD_600_ was transferred into a 1.5 mL microcentrifuge tube. Cells were then centrifuged for 10 min at 13,000 rpm and 4°C. The supernatant was discarded and cell pellets were resuspended in 1 mL of ice-cold phosphate buffer saline (PBS, Cat#10722497, Invitrogen). Centrifugation and PBS washing was then repeated. The supernatant was discarded and cell pellets were stored at -80°C for 1 h. Finally, pellets were resuspended in 100 μL of ammonium carbonate ((NH_4_)_2_CO_3_ (Cat#10361-29-2, Honeywell), and centrifuged for 10 min at 4°C. The supernatant was stored at -80°C until analysis by FI-MS.

### Flow-injection mass spectrometry (FI-MS)

For FI-MS cell extracts were directly injected into an Agilent 6546 Series quadrupole time-of-flight mass spectrometer (Agilent Technologies, USA) as described previously^9,10^. The electrospray source was operated in negative and positive ionization mode. The mobile phase was 60:40 isopropanol:water buffered with 10 mM ammonium carbonate (NH_4_)_2_CO_3_ and 0.04 % (v/v) ammonium hydroxide for both ionization modes, and the flow rate was 0.15 mL/min. For online mass axis correction, 2-propanol (in the mobile phase) and HP-921 were used for negative mode and purine and HP-921 were used for positive mode. Mass spectra were recorded from 50 to 1700 m/z with a frequency of 1.4 spectra/s for 0.5 min using 10 Ghz resolving power. Source temperature was set to 225°C, with 1 L/min drying gas and a nebulizer pressure of 20 psi. Fragmentor, skimmer, and octupole voltages were set to 120 V, 65 V, and 650 V, respectively. Capillary voltage was set to 3,500 V. For FI-MS of whole living cells, 3 μL of an *E. coli* culture in 1:8 M9-medium:water was injected. The heat-exchanger was set to 90°C, and capillary voltage was 5,000 V.

Raw data files were converted into mzXML files and processed by custom MATLAB scripts. The 32 spectra with the highest signal in the total ion count were summed and baseline adjusted with *msbackadj*.*m*. Peaks with a minimum peak height of 5000 units and a peak prominence of 5000 units were selected with *findpeaks*.*m*, and annotated with a 3 mDa tolerance by matching monoisotopic masses of all metabolites in the iML1515 model^13^, considering a single proton loss ([M-H]-) in negative mode and single proton gain ([M+H]+) in positive mode. Positive and negative mode annotation were merged and if a metabolite was annotated in both modes negative mode was selected. For each metabolite, the height of the annotated ion peak was taken for further analysis and normalized to the mean of the control strain to obtain fold-changes.

For barcode detection raw Agilent files were converted into ‘‘mzML’’ format using MSConvert and mzML files were used for FLASHDeconv to obtain monoisotopic masses.

